# Phyllosphere microbes decouple nutrient enrichment from plant productivity

**DOI:** 10.64898/2026.08.16.745140

**Authors:** Na Wei

**Author notes:** **Author for correspondence:** Na Wei.

## Abstract

Resource limitation theory predicts that nutrient enrichment enhances plant productivity. Yet plant-associated microbes can modify this relationship by facilitating nutrient acquisition, competing for resources, or restructuring the plant–environment interface. These processes generate contrasting predictions for whether added nutrients are converted into plant population growth. Whether phyllosphere microbiomes mediate this resource–productivity relationship remains unclear. Here we show that phyllosphere microbiomes decoupled nutrient enrichment from productivity in a duckweed polyploid complex. In microbe-free microcosm ecosystems, nutrient enrichment increased productivity, whereas with microbes, enrichment failed to increase productivity despite abundant residual nutrients. This decoupling was not explained by direct microbial competition for nutrients or predicted microbial functions associated with pathogenicity, oxygen depletion, or acidification. Instead, nutrient enrichment stimulated biofilm formation, potentially restricting plant access to nutrients. Predicted microbial phosphorus immobilization and transformation also increased, but neither explained the decline in ecosystem phosphorus removal. This decline instead reflected lower plant productivity associated with biofilm formation, leaving much of the phosphorus unused. These patterns were consistent across ploidy levels, with the productivity advantage of polyploids associated with greater nutrient-use efficiency rather than greater tolerance of microbial effects. Our results reveal that phyllosphere microbiomes mediate how nutrient enrichment translates into ecosystem functioning.

## Introduction

The global prevalence of nutrient enrichment across terrestrial and aquatic ecosystems is rapidly altering ecosystem productivity and other key functions [1–5]. Resource limitation theory predicts that enrichment of biologically limiting nutrients can support larger population sizes and higher standing biomass [6, 7]. Resource limitation arises when nutrient availability falls short of biological demand. Enrichment is therefore expected to increase productivity when it alleviates this constraint, unless it simultaneously strengthens other abiotic or biotic constraints on population growth. Although sustained enrichment can also reduce productivity indirectly by eroding plant diversity [1–3, 8, 9], direct resource-driven gains generally outweigh these diversity-mediated declines [4, 5]. Beyond the diversity-mediated effects, plant-associated microbiomes may provide an additional pathway through which enrichment alters productivity, even when plant diversity remains unchanged. Yet whether microbial symbionts reinforce, counteract, or decouple the expected positive effect of nutrient enrichment on plant productivity remains unclear.

Microbial communities are important regulators of plant population dynamics and ecosystem productivity [10, 11]. Nutrient enrichment can alter microbial richness, community structure, and function. These responses include shifts microbial life-history strategies, functional potential, and the balance between mutualistic and antagonistic taxa [5, 12–16]. However, most work has focused on how enrichment reshapes microbial communities themselves rather than on whether these changes modify the effects of enrichment on plant productivity. Evidence linking nutrient enrichment to microbially mediated changes in ecosystem functioning remains concentrated in terrestrial soil microbiomes and has primarily concerned soil multifunctionality [14, 15]. We have far less understanding of these microbial responses in the phyllosphere, an important interface between plants and their surrounding environment. As a result, whether nutrient-induced changes in phyllosphere microbiomes feed back to alter plant productivity remains poorly understood.

Plant-associated microbes can influence plant productivity through multiple mechanisms that enhance or suppress plant population growth, depending on environmental context. Microbes can increase nutrient availability by mineralizing organic compounds, altering redox states, or transforming resources into more plant-accessible forms [17–19]. At the same time, they may compete with plants for nutrients or sequester them in microbial biomass, thereby reducing the pool available for plant uptake [20–23]. Phosphorus exemplifies this duality: microbial communities can increase plant access to phosphorus through mineralization and solubilization but can also immobilize it and limit its bioavailability [22, 24]. Nutrient enrichment can further shift plant–microbe interactions along the mutualism–parasitism continuum, weakening nutrient-provisioning interactions or favoring antagonistic strategies, as seen in rhizobia, mycorrhizal fungi, and other microbial guilds [25–28]. Microbes may also suppress productivity by imposing biotic stress or modifying the physicochemical environment at plant surfaces. Virulence factors can directly impair plant performance, whereas biofilms can restructure the plant–environment interface. Although biofilms may stabilize microbial assemblages and buffer environmental fluctuations, they can also restrict nutrient diffusion, retain resources within the extracellular matrix, and generate localized oxygen depletion or acidification [17, 29–32]. Because these pathways can act synergistically or in opposing directions and vary with nutrient availability, the net effect of microbial communities on plant productivity under enrichment remains difficult to predict.

The effects of nutrient enrichment on plant–microbe symbiosis and productivity may also vary among hosts. Host variation can influence plant resource demand, stress sensitivity, and the assembly of associated microbiomes. Polyploidy is a particularly important axis of host variation. Whole-genome duplication (WGD) has recurred throughout plant evolution [33, 34]. WGD is frequently associated with enhanced tolerance to resource limitation and other abiotic stresses, potentially contributing to broader ecological amplitudes and higher fitness in stressful environments, although these effects vary among lineages and environments [35–38]. WGD also alters morphology, physiology, and metabolic demand [39–41], which may reshape microbial colonization and composition [42, 43]. Together, these differences raise the possibility that WGD may alter the net microbial influence on plant productivity under nutrient enrichment, by shifting the balance among nutrient provisioning, nutrient competition, and stress buffering.

Floating aquatic plants such as duckweeds (Lemnaceae) offer an exceptional system [11, 44–47] for determining how phyllosphere microbiomes mediate the magnitude and direction of nutrient enrichment effects on plant productivity and ecosystem functioning. Because duckweeds possess a highly reduced body plan [44, 48], the frond–water interface (the phyllosphere) provides a major site of plant–microbe interactions. Their rapid growth, short generation times, and nutrient-limited population dynamics [48] facilitate the experimental quantification of nutrient-driven changes in plant productivity. WGD has also recurred frequently in duckweed populations, particularly in *Lemna* and *Wolffia* [49]. To isolate the effects of WGD from other variation among hosts, we generated clonal synthetic autotetraploids for comparison with their diploid progenitors. We replicated this ploidy comparison across two of the most widespread duckweed species, *Lemna minor* and *Lemna japonica*. Here, using a microcosm experiment that manipulated nutrient supply, microbial presence, and ploidy level, we ask: (1) whether microbiomes alter the productivity gain from nutrient enrichment predicted by resource limitation theory, and whether this effect depends on ploidy; (2) how these factors influence ecosystem phosphorus removal; and (3) which microbial functional pathways are associated with the reinforcement, suppression, or decoupling of nutrient enrichment effects on plant productivity and phosphorus removal. Together, these tests aimed to improve our understanding of how microbiomes and host ploidy shape the fundamental links between resource limitation, plant productivity, and ecosystem functioning.

## Methods

### Duckweed genotypes and synthetic polyploids

We focused on six diploid (2*x*) duckweed genotypes, including two *Lemna minor* and four *Lemna japonica*. These genotypes were collected in Ohio, USA. To generate synthetic autotetraploids (4*x*), approximately 120 individuals of each axenic genotype were grown in 15 mL of 0.5× Hoagland salts supplemented with 0.5% sucrose and treated with 100 µM oryzalin with DMSO (0.33 mL L^-1^). Treated individuals were kept overnight in darkness and then grown at 24 °C under 16-h days for one to two weeks, depending on the genotype. Surviving individuals were rinsed twice with sterile water and propagated individually for six months before ploidy screening to identify stable synthetic 4*x* lines. Ploidy was examined using flow cytometry following earlier work [40]. For each original genotype, we established a lineage comprising a synthetic 4*x* line, a treated but unconverted 2*x* line (2*x*.nc), and the untreated wild 2*x* line. The 2*x*.nc lines controlled for exposure to oryzalin and DMSO.

### Nutrient enrichment experiment

The experiment used a three-way factorial design that manipulated ploidy level (2*x*, 2*x*.nc, and 4*x*), nutrient level (low: 0.025× Hoagland salts with 0.0125% sucrose; high: 0.1× Hoagland salts with 0.05% sucrose), and microbiome treatment (M0: absent; M1: present). Both nutrient levels were within the natural range experienced by wild duckweeds [50]. The design was replicated four times, yielding 288 microcosms: 6 lineages × 3 ploidy levels × 2 nutrient levels × 2 microbiome treatments × 4 replicates.

At the start of the experiment, six axenic duckweed individuals were introduced into each microcosm containing 20 mL of sterile medium. Microbiome treatments were applied two days later. Microbial inoculants were obtained from pond water collected from a local duckweed habitat in the Cleveland area using two sterile 50 mL centrifuge tubes. Microbes were pelleted by centrifugation at 4400 rpm for 10 min, resuspended in 20 mL of 0.25× phosphate buffered saline (PBS), and passed through a 1.2 μm sterile PES syringe filter to remove eukaryotic cells. Each M1 microcosm received 100 µL of microbiome extract, whereas each M0 microcosm received 100 µL of sterile 0.25× PBS. Microcosms were randomized and rotated daily in a growth room maintained at 24 °C under a 16-h photoperiod. The experiment lasted two weeks, by which time population growth had reached equilibrium.

At the end of the experiment, duckweeds were harvested and sonicated at 40 kHz for 5 min to recover duckweed-associated microbes. Microbes were pelleted by centrifugation at 4400 rpm for 10 min and stored at -20 °C until DNA extraction. To measure ecosystem phosphorus removal, 13 mL of liquid from each microcosm was centrifuged at 4400 rpm for 10 min. The supernatant was used to measure residual total phosphorus, including inorganic and organic forms [11]. Plant productivity was quantified as the dry biomass of harvested duckweeds after drying at 60 °C for 72 h.

### Duckweed bacterial community sequencing

Microbial DNA was extracted using a modified cetyltrimethylammonium bromide (CTAB) protocol [50]. DNA samples were sent to Argonne National Laboratory for library preparation targeting the bacterial 16S rRNA V5–V6 region (799F: AACMGGATTAGATACCCKG; 1115R: AGGGTTGCGCTCGTTG). Sequencing was performed on an Illumina MiSeq platform using 250 bp paired-end reads.

Paired-end reads were processed using DADA2 v1.32.0 [51] in R v4.4.1 [52] following previous pipelines [53, 54]. Reads were quality filtered [truncLen=c(200, 200), trimLeft=c(30, 10), maxN=0, truncQ=2, rm.phix=T, maxEE=c(2,2)] prior to amplicon sequence variant (ASV) inference and chimera removal. ASVs were taxonomically assigned using the SILVA NR99 reference database (release 132) implemented in DADA2. The resulting ASV table was filtered to remove non-bacterial ASVs. Samples were rarefied to 10,000 reads (median depth before rarefaction: 30,737 reads; Fig. S1). Five samples had fewer than 10,000 reads (one with 4,416 reads and four with more than 8,000 reads), but showed saturated rarefaction curves. These samples were retained and were proportionally normalized to 10,000 reads [50]. Rare ASVs (< 0.001% of total observations) were removed. The final dataset contained 864 ASVs across 143 duckweed microbiome samples from M1 microcosms; one M1 sample failed during sequencing.

### Statistical analyses of plant productivity and residual phosphorus

To test whether microbiomes modified the effect of nutrient enrichment on plant productivity across ploidy levels, we fitted a linear mixed model (LMM) using lme4 [55]. Ploidy level, nutrient level, microbiome treatment, and their two-and three-way interactions were included as fixed effects. Species and lineage nested within species were included as random effects. We used the same model structure to analyze ecosystem phosphorus removal based on residual phosphorus (OD885). As the model containing both random effects failed to converge, we retained species as the only random effect because it explained more variation than lineage. Response variables were power-transformed when necessary, with optimal parameters determined using the Box–Cox method. Plant productivity required no transformation, whereas residual phosphorus was log-transformed. Statistical significance (type III sums of squares), least-squares means and contrasts were evaluated using lmerTest [56] and emmeans [57].

### Statistical analyses of predicted microbial functional potentials

To test candidate mechanisms through which microbiomes diminished plant productivity under nutrient enrichment, we quantified predicted functional potentials associated with biotic and abiotic stress. These included pathogenesis, biofilm formation, micro-oxia, and acidification. Microbial functional potentials were inferred using PICRUSt2 [58]. The resulting KEGG Orthology (KO) abundance matrix was converted to relative abundances. To address the constant-sum constraint of compositional data, KO relative abundances were centered log-ratio (CLR) transformed using compositions [59], with zeros first imputed using Bayesian-multiplicative replacement in zCompositions [60]. KO identifiers were annotated using the KEGG database (https://rest.kegg.jp/list/ko; accessed October 21, 2025).

Pathogenic functional potential was evaluated using KOs representing the Type III Secretion System (T3SS), a conserved injectisome used to deliver host-manipulating effectors [61, 62], and KOs encoding cell wall degrading enzymes (CWDE) [63, 64], which mediate plant cell wall breakdown and indicate potential pathogenic or saprotrophic capacities (Table S1). Biofilm functional potential was assessed using core KOs involved in exopolysaccharide synthesis, quorum sensing, flagellar motility, and chemotaxis. These processes coordinate surface approach, attachment, and matrix formation (Table S1). Micro-oxic functional potential was evaluated using KOs associated with aerobic respiration and oxygen-scavenging pathways, which may contribute to localized oxygen depletion and micro-oxic boundary layers (Table S1). Acidification functional potential was quantified using KOs linked to organic acid production and fermentation, which can lower local pH (Table S1).

To test candidate microbial processes influencing residual phosphorus, we quantified KOs associated with phosphorus immobilization and transformation. Phosphorus immobilization functional potential was evaluated using KOs associated with high-affinity phosphate uptake (‘scavenge mode’) [65, 66] and polyphosphate synthesis and storage (‘storage mode’) [67, 68]. These functions indicate the potential to sequester phosphorus in microbial biomass or intracellular polyphosphate granules. Phosphorus transformation functional potential was quantified using KOs involved in phosphorus turnover. These included alkaline phosphatases, phosphonate-degradation pathways, and other enzymes involved in phosphorus release or the cleavage of phosphorus-containing compounds within microcosms.

Functional potential indices were calculated as the mean CLR-transformed values of the KOs assigned to each functional group. Because functional profiles were available only for M1 microcosms, we fitted linear models (LMs) with ploidy level, nutrient level, and their interaction as predictors, and functional potentials indices as response variables. Random effects for species and lineages were not included due to their non-significant effects and model convergence challenge.

### Structural equation modeling (SEM)

To evaluate how nutrient enrichment influenced microbial functional potentials, plant productivity, and microcosm residual phosphorus, we fitted SEM using lavaan [69]. The analysis included only M1 microcosms, for which microbial functional profiles were available. Nutrient treatment was coded as a binary variable representing low (0) and high (1) nutrient levels, such that path coefficients represented the effects of nutrient enrichment. Ploidy level was not included as an exogenous predictor because its effects on microbial functional potentials and residual phosphorus were negligible. Variable transformations followed those used in the LMMs (no transformation for plant productivity; log transformation for residual phosphorus). The initial full SEM (Fig. S2) was fitted using maximum likelihood with robust Huber–White standard errors. To reduce model complexity, we re-fitted the model retaining statistically supported paths (*P* ≤ 0.05). Model fit was confirmed: comparative fit index (CFI > 0.90), root mean square error of approximation (RMSEA; lower bound of the 90% CI < 0.05), and standardized root mean square residual (SRMR < 0.10).

## Results

In the absence of microbes (M0), nutrient enrichment consistently boosted plant productivity across ploidy levels, with high-nutrient microcosms producing 33–55% more dry biomass than low-nutrient microcosms (all *P* < 0.001; Fig. 1a). This positive nutrient response was lost once microbes were introduced. Under M1, nutrient enrichment no longer increased plant productivity (2*x*, *P* = 0.4; 2*x*.nc, *P* = 0.2; 4*x*, *P* = 0.044; Fig. 1a). Across both nutrient levels, plants grown with microbes were consistently less productive than microbe-free plants (all *P* < 0.05).

**Figure 1.**
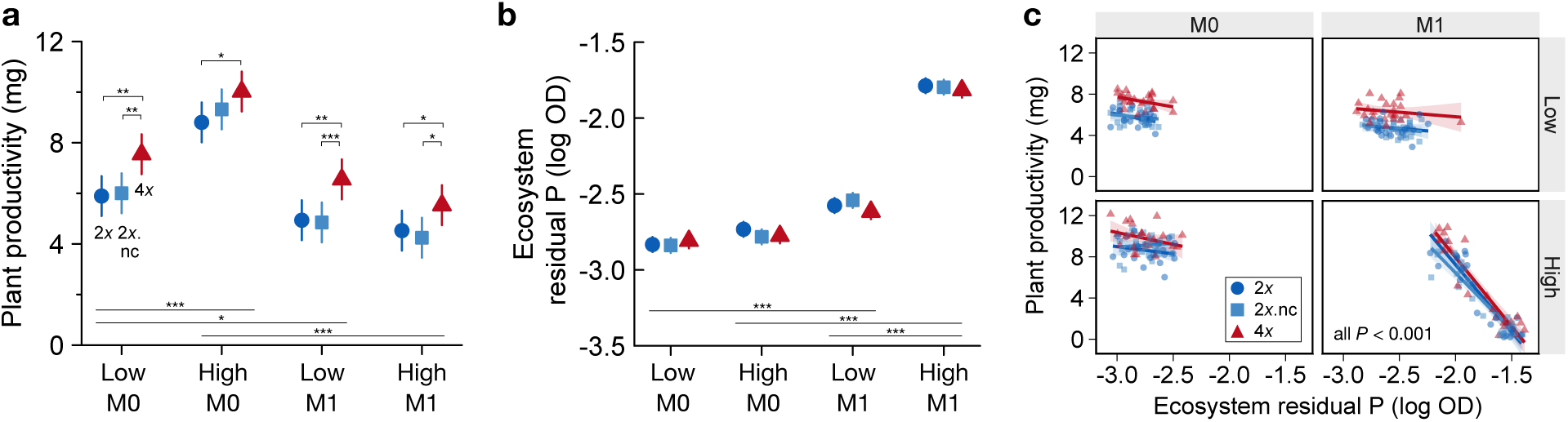
Microbial communities decouple nutrient enrichment from plant productivity. (a) Plant productivity, measured as dry biomass, across ploidy levels (2*x*, wild diploids; 2*x*.nc treated but unconverted diploids; 4*x*, synthetic autotetraploids) under low and high nutrient supply in microbe-free (M0) and microbe-present (M1) microcosms. (b) Residual phosphorus, measured as absorbance at 885 nm (OD885), at the end of the experiment. Points in (a) and (b) show least-squares means (± SE) estimated from linear mixed models (predictors: ploidy level, nutrient level, microbiome treatment, and all two-way and three-way interactions; *n* = 288). Significant contrasts within or between treatments are indicated: \*\*\**P* < 0.001; ** *P* < 0.01; \**P* < 0.05. (c) Relationship between plant productivity and residual phosphorus. Lines show linear model fits with shaded 95% confidence intervals.

One explanation is that microbes might compete with plants for nutrients, thereby intensifying plant nutrient limitation. To test this, we quantified residual total phosphorus in microcosms (Fig. 1b). If microbial competition had directly depleted phosphorus, microcosms with microbes, especially under high nutrient supply, should have shown lower residual phosphorus. Instead, the opposite pattern emerged: residual phosphorus was substantially higher in microcosms with microbes (all *P* < 0.001; Fig. 1b).

In microbe-free microcosms, plants removed most of the supplied phosphorus, leaving consistently low residual phosphorus across nutrient levels (Fig. 1b,c) and revealing a tight coupling between nutrient supply and plant productivity. Introducing microbes disrupted this coupling. Although plant productivity remained positively related to phosphorus uptake (Fig. 1c; all *P* < 0.001), a large fraction of the added phosphorus was not assimilated by plants when microbes were present. Thus, productivity suppression was inconsistent with phosphorus depletion through direct microbial competition and instead pointed to reduced plant phosphorus acquisition despite phosphorus remaining in microcosms.

Genome duplication did not alter this microbial decoupling of nutrient enrichment from plant productivity. Polyploids consistently achieved higher productivity than diploids across treatments (Fig. 1a). Yet residual phosphorus did not differ among ploidy levels (Fig. 1b). This pattern suggested that the polyploid advantage arose through mechanisms other than enhanced phosphorus acquisition, potentially including greater nutrient-use efficiency or resistance to microbe-induced stress (which was ruled out later).

To examine whether predicted microbial functions associated with biotic stress could explain reduced productivity, we tested whether pathogenic functional potential increased under high nutrient supply (Fig. 2a). Contrary to this expectation, pathogenic functional potential was higher under low nutrient supply across ploidy levels (all *P* < 0.001, Fig. 2a). This pattern was consistent for both T3SS and CWDE components (Fig. S3), indicating that predicted pathogenic or saprotrophic capacity in the phyllosphere was higher when external resources were limited. However, pathogenic functional potential showed no statistically supported relationship with plant productivity in the SEM (Fig. 3).

**Figure 2.**
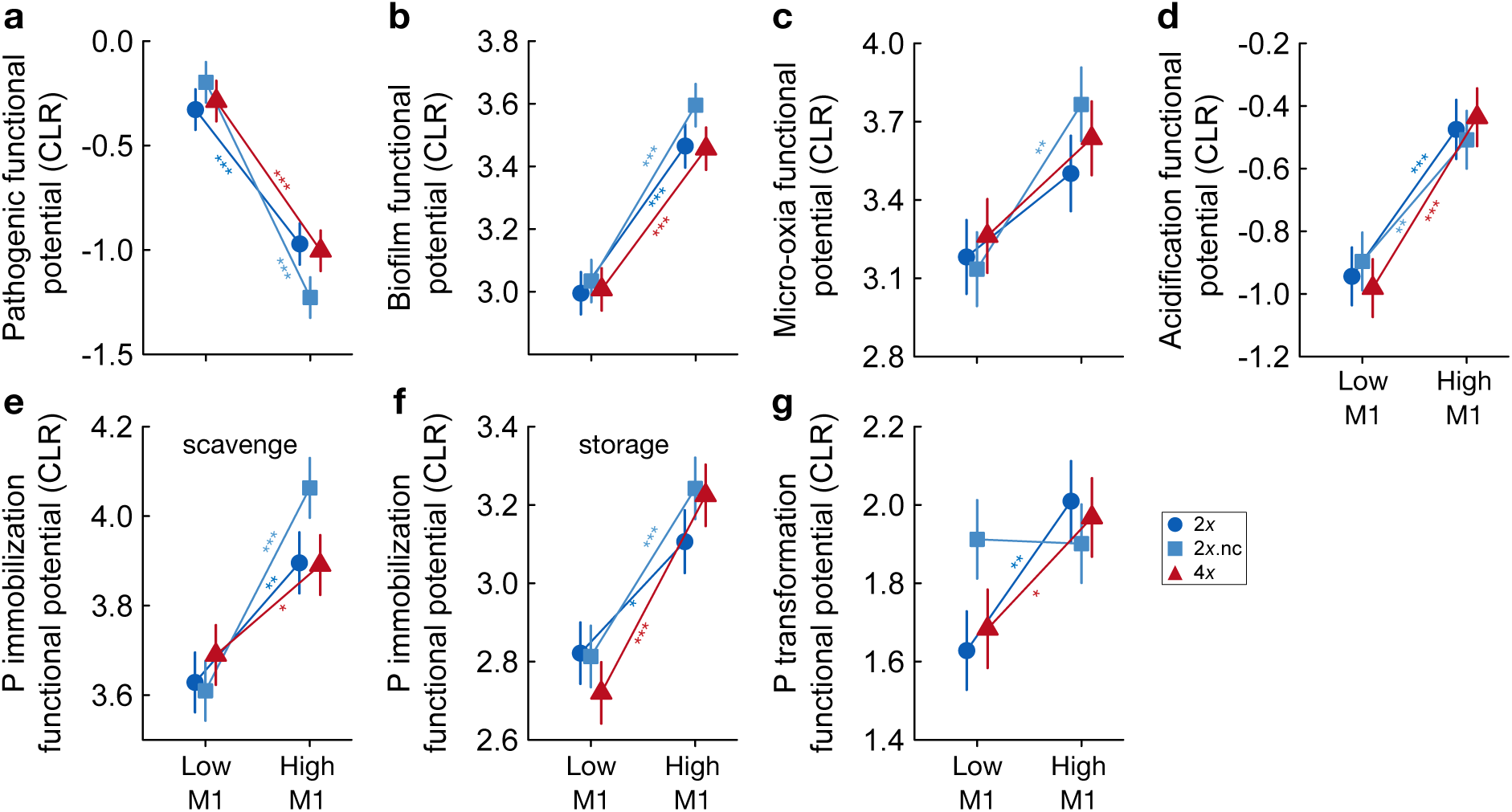
Nutrient enrichment alters predicted microbial functional potential. (a–g) Predicted microbial functional potentials, quantified as the mean CLR-transformed relative abundances of KEGG Orthologs assigned to each functional category, across nutrient levels (low and high) and ploidy levels (2*x*, wild diploids; 2*x*.nc treated but unconverted diploids; 4*x*, synthetic autotetraploids) in microbe-present microcosms (M1). Points show least-squares means (± SE) estimated from linear models (predictors: ploidy level, nutrient level, and their two-way interaction; *n* = 143). Significant contrasts between nutrient levels are indicated: \*\*\**P* < 0.001; ** *P* < 0.01; \**P* < 0.05.

**Figure 3.**
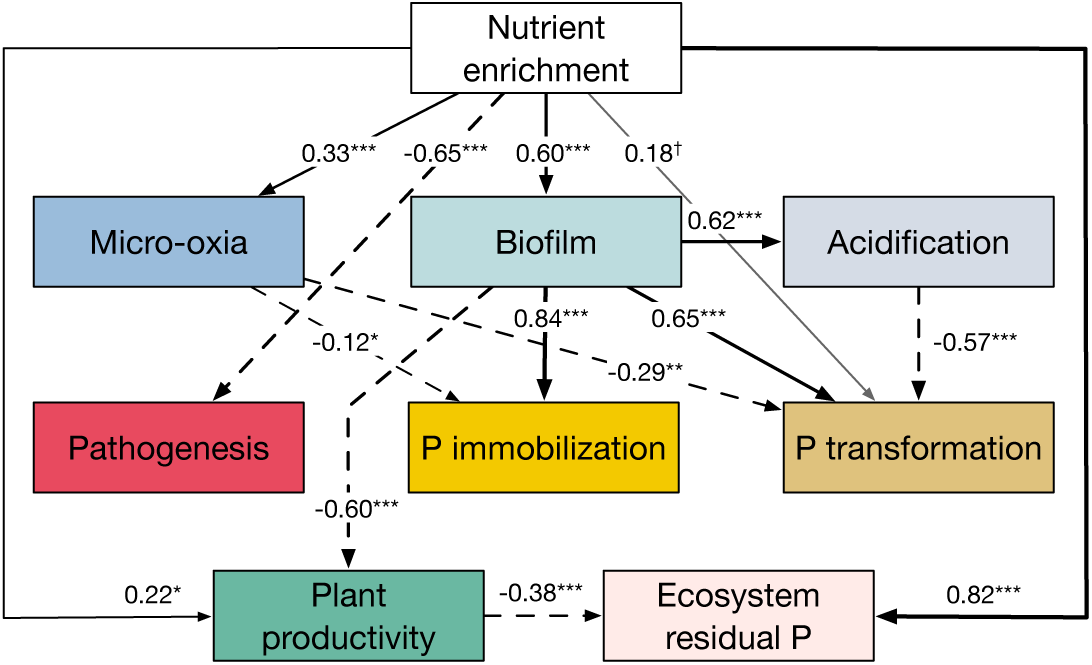
A microbially driven pathway decouples nutrient supply from plant productivity. The reduced structural equation model (SEM) describes how nutrient enrichment alters predicted microbial functional potentials and how these changes influence plant productivity and residual phosphorus. Black arrows indicate significant positive (solid) and negative (dashed) pathways; grey arrows indicate nonsignificant pathways. Numbers adjacent to arrows are standardized path coefficients. The SEM was fitted using maximum likelihood with robust Huber–White standard errors. Nutrient treatment was coded with the low-nutrient level as the reference category; coefficients therefore represent the effects of nutrient enrichment. Predicted microbial functional potentials were quantified as the mean CLR-transformed relative abundances of KEGG Orthologs assigned to pathogenesis, micro-oxia, biofilm formation, acidification, phosphorus immobilization (combining scavenging and storage modes), and phosphorus transformation. Significance levels: \*\*\**P* < 0.001; \*\**P* < 0.01; \**P* < 0.05; grey arrow, *P* > 0.05. The full SEM is presented in Figure S2.

We next evaluated predicted microbial functional potentials that could generate physical or abiotic constraints. Biofilm functional potential increased under nutrient enrichment (all *P* < 0.001; Fig. 2b), consistent with visible biofilm development in the phyllosphere. Biofilm functional potential emerged as a major driver of reduced plant productivity (*r* = -0.60, *P* < 0.001; Fig. 3). Micro-oxia and acidification functional potentials were also elevated under nutrient enrichment (Fig. 2c,d), yet neither had a notable effect on plant productivity (Fig. 3).

Lastly, we examined whether predicted microbial potentials for phosphorus immobilization and transformation could explain residual phosphorus. For immobilization, we expected the scavenge mode to predominate under low nutrient supply and the storage mode under high nutrient supply. Instead, both modes increased under nutrient enrichment in the phyllosphere (all *P* < 0.05; Fig. 2e,f). Biofilm functional potential was strongly positively related to both modes (*r* = 0.84, *P* < 0.001; *r* = 0.65, *P* < 0.001; Fig. 3). However, neither phosphorus immobilization nor transformation functional potential was related to residual phosphorus in microcosms (Fig. 3). Instead, lower plant productivity led to higher residual phosphorus (*r* = -0.38, *P* < 0.001; Fig. 3). This relationship may reflect lower plant phosphorus uptake associated with biofilm-related productivity suppression, leaving more of the supplied phosphorus unused.

## Discussion

Our experiment showed that whether nutrient enrichment increased plant productivity, as predicted by resource limitation theory, depended on microbial context. Without microbes, enrichment increased productivity; with microbes, this response disappeared. This decoupling was not explained by direct nutrient competition, predicted pathogenicity, micro-oxia, or acidification. Instead, enrichment promoted biofilm formation, which emerged as the leading candidate mechanism restricting nutrient acquisition at the plant–water interface. Phyllosphere microbes thus acted as ecosystem engineers, reshaping access to resources rather than simply depleting them. Polyploidy increased plant productivity but did not alter this microbial control. Together, the productivity benefit of enrichment was not an intrinsic property of nutrient supply, but an emergent property shaped by phyllosphere microbiomes.

### Biofilms may impose microscale barriers

The association between biofilm formation and nutrient–productivity decoupling is consistent with principles of microbial spatial ecology. Biofilms can create chemical and diffusive gradients at biological surfaces by thickening boundary layers and accumulating extracellular polymeric substances (EPS) [17, 29–32]. Nutrient enrichment can promote this surface-attached growth through increased EPS production, quorum sensing, and adhesion [30, 32, 70]. Depending on context, the resulting matrix may retain beneficial metabolites and buffer environmental stress [18, 71, 72]. However, it may also impede nutrient diffusion or sequester resources before they reach plant surfaces [30, 32].

Biofilms can also generate micro-oxic or acidic microsites through restricted diffusion and microbial metabolism [30, 32]. In our experiment, biofilm potential increased acidification but not micro-oxia potential (Fig. 3), and neither explained reduced plant productivity. Biofilm formation instead emerged as the leading candidate mechanism underlying nutrient–productivity decoupling. By accumulating at the plant–water interface, biofilms may have restricted plant access to nutrient even while it remained abundant in the surrounding environment. This mechanism is particularly relevant in *Lemna*, where root contribution was minimal and nutrient exchange occurred primarily across the frond–water interface. Phyllosphere biofilms may therefore generate resource limitation at the microscale by controlling access to otherwise abundant resources.

### Other predicted functional shifts exert limited effects

Nutrient enrichment altered multiple predicted microbial functions, but these changes did not explain nutrient–productivity decoupling. Pathogenesis-related functional potential was higher under low, not high, nutrient supply. Under nutrient scarcity, microbes may rely more strongly on traits that release limiting resources from host-derived or other organic substrates. Many functions classified as pathogenic, including cell wall-degrading enzymes (CWDEs), can also support surface colonization, opportunistic nutrient acquisition, or saprotrophy without causing active infection [63, 64, 73]. This functional versatility highlights the context dependence of microbial function: the same function can serve different ecological roles under different conditions. Thus, elevated predicted pathogenic potential may not translate into substantial plant damage.

Abiotic stress pathways showed a similar disconnect between microbial responses and plant productivity. Enrichment directly increased predicted micro-oxia potential, whereas its effect on acidification potential was mediated through biofilm formation. The direct response may reflect enrichment-induced increases in microbial respiration and oxygen demand, whereas the indirect pathway may arise because biofilms concentrate fermentative and acidifying activity within their surface-attached matrix [15, 70, 74–76]. These predicted functional shifts captured the expected microbial responses to enrichment. But neither generated sufficient stress to limit plant productivity under our experimental conditions. Their effects on productivity may have been buffered by the surrounding water or remained below the plants’ tolerance thresholds.

### Plant productivity governs ecosystem phosphorus removal

Biofilms substantially reshaped microbial phosphorus immobilization and transformation. We initially expected a clear separation between the scavenge mode of phosphorus immobilization, characterized by high-affinity uptake under low phosphorus levels [65, 66], and the storage mode, characterized by polyphosphate accumulation under high phosphorus levels [67, 68, 77]. Instead, both modes increased under nutrient enrichment. This simultaneous rise might be caused by the microscale environment created by enrichment-induced biofilms. Diffusion limitation within the EPS matrix may have generated localized phosphorus depletion, activating high-affinity scavenging pathways, while the high external phosphorus availability simultaneously enabled polyphosphate storage. Biofilms also elevated phosphorus transformation potential, consistent with dense microbial aggregates stimulating enzymatic turnover of organic phosphorus [78]. In contrast, predicted micro-oxia potential was negatively associated with both phosphorus immobilization and transformation potentials, contrary to reports that low-oxygen microzones can promote phosphatase activity or phosphonate degradation [79–82]. This pattern suggests that the functional consequences of micro-oxia may depend on the intensity and spatial extent of oxygen limitation. Acidification similarly reduced transformation potential, consistent with the pH sensitivity of alkaline phosphatases that lose activity under acidic conditions.

Neither immobilization nor transformation explained variation in residual phosphorus. Instead, residual phosphorus accumulated because biofilm-mediated suppression of plant productivity left much of the supplied phosphorus unused. This is consistent with our previous finding that duckweed population size, rather than microbial communities, was the primary driver of ecosystem phosphorus removal [11].

### WGD enhances productivity but not microbial mediation

Resource limitation depends not only on nutrient supply but also on host traits. Whole-genome duplication provided a tractable test of whether host variation alters phyllosphere microbiome functional potentials and thereby modifies the translation of nutrient enrichment into plant productivity.

Consistent with widespread evidence of polyploidy advantage [36–38], synthetic tetraploids maintained higher productivity than their diploid counterparts across treatments. This advantage was not associated with greater phosphorus acquisition, as residual phosphorus did not differ among ploidy levels. Instead, the greater productivity of synthetic tetraploids at similar residual phosphorus levels is consistent with more efficient use of acquired nutrients.

Importantly, however, WGD did not alter the direction or magnitude of microbial mediation: polyploids experienced the same microbially driven decoupling of the nutrient– productivity relationship as diploids. Predicted functional potentials of pathogenesis, abiotic stress, and phosphorus cycling also responded similarly across ploidy levels. Although polyploids often assemble microbial communities that differ compositionally from those of their diploid progenitors [42, 43], the functional consequences remain uncertain. Functional redundancy may allow compositionally distinct microbiomes to retain similar functional capacities [83–85]. Such redundancy may help explain why microbiome functional responses to nutrient enrichment remained similar across ploidy levels despite shifts in individual ASVs.

Whether wild polyploids recruit microbial partners with sufficiently distinct functional roles to modify this outcome remains an open question.

In summary, our findings highlight a microbial mechanism underlying nutrient– productivity decoupling. Phyllosphere microbiomes may reshape the interface between resource supply and plant uptake. They may thereby determine whether enrichment alleviates resource limitation or creates new barriers to resource acquisition. The consequences can extend from plant productivity to ecosystem phosphorus removal. Whether such microbial mediation operates in natural ecosystems will depend on other processes such as hydrodynamics, nutrient flows, and microbial interactions. These factors may shape both biofilm dynamics and the broader functional effects of microbiomes. Moreover, because nutrient fluxes at plant surfaces were not measured and microbial functions were inferred from taxonomic marker data, the proposed mechanism remains to be tested directly. Future work should therefore establish its causal basis and determine the range of conditions under which it operates. Resource limitation is therefore not determined solely by ecosystem resource supply but can be an emergent property shaped by phyllosphere microbiomes.

### Ethics

This work did not require ethical approval from a human subject or animal welfare committee.

## Data accessibility

All data that support the findings of this study are included in this published article, its electronic supplementary material, and are available at Science Data Bank (ScienceDB; https://www.scidb.cn/s/6zUZ7n)

## Declaration of AI use

I have not used AI-assisted technologies in creating this article.

## Authors’ contributions

N.W.: Conceptualization, Data curation, Formal analysis, Funding acquisition, Investigation, Methodology, Resources, Visualization, Writing – original draft, Writing – review & editing

## Acknowledgements.

I thank Maris Hollowell and Emily Lewis for assistance with the experiments and James Gossard for assistance with ploidy screening.

---

**Figure S1.**
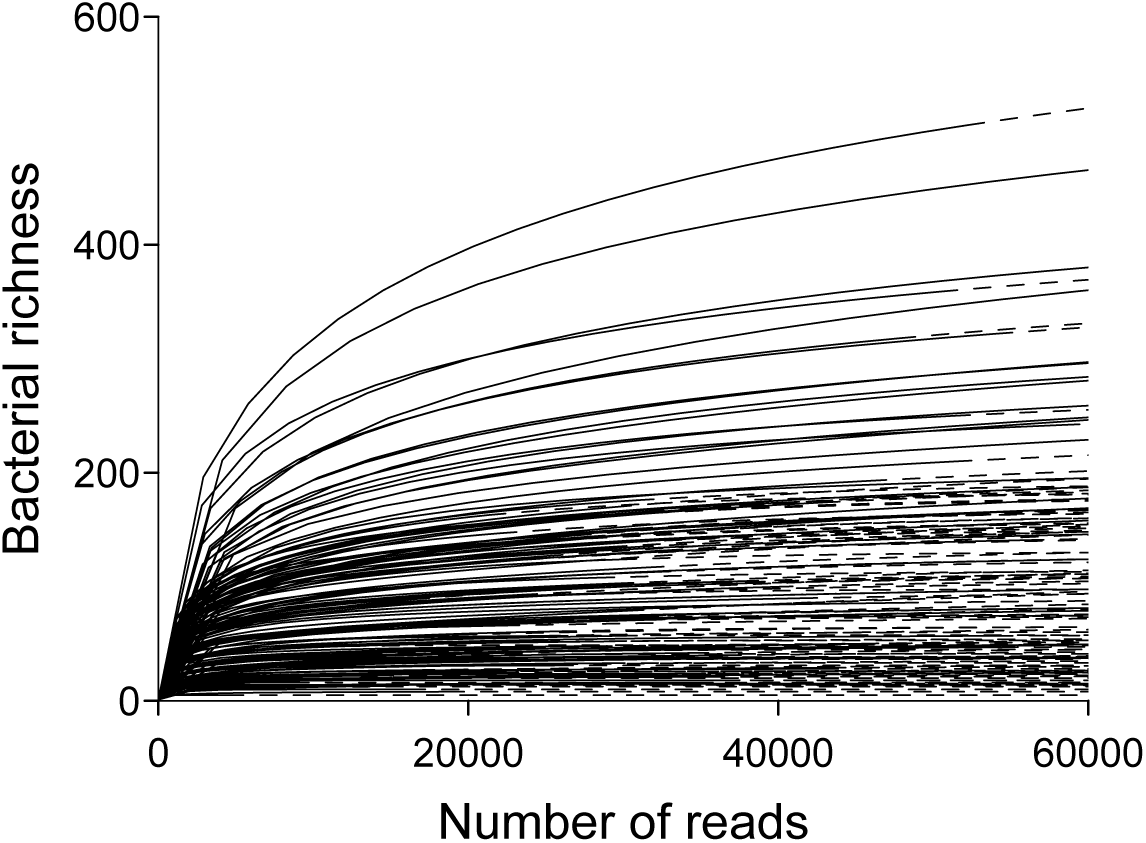
Rarefaction shows that the majority of bacterial richness was captured with our sequencing effort. The number of reads (*x*-axis) obtained for bacterial microbiomes in individual experimental microcosms is represented by the solid portion of each line, whereas the dashed portion indicates extrapolation in the rarefaction analysis using the R package iNEXT.

**Figure S2.**
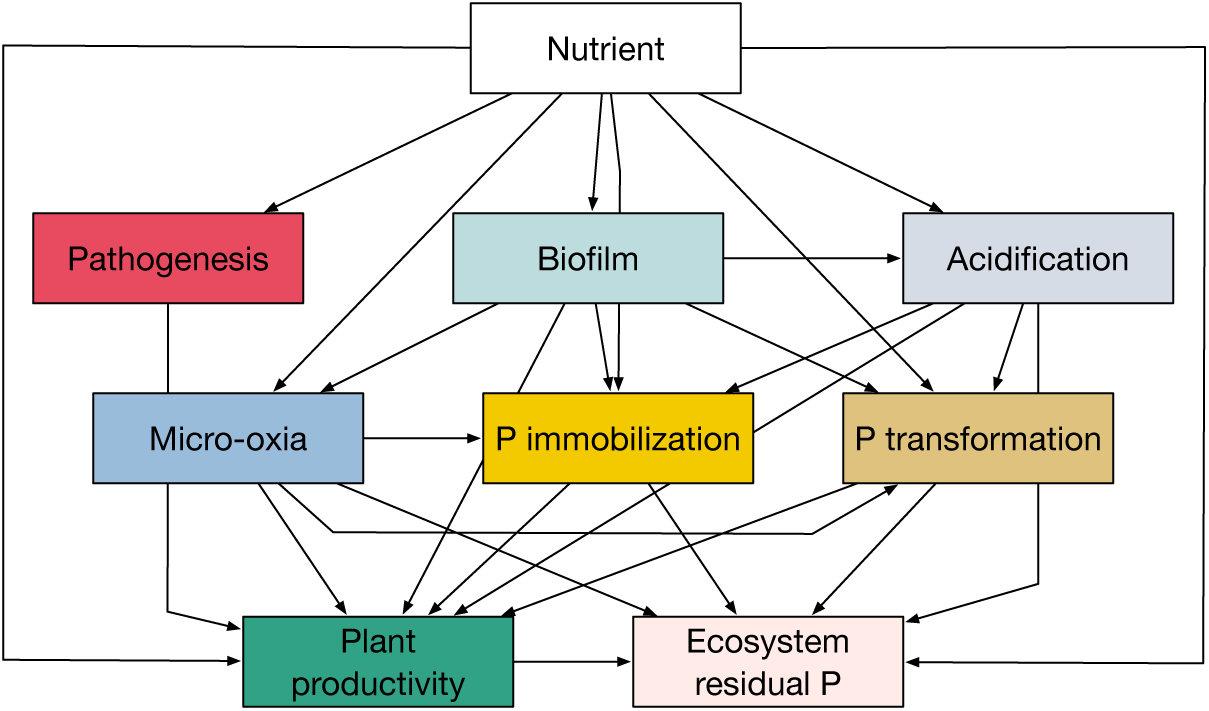
Full structural equation model (SEM) depicting all hypothesized paths linking nutrient enrichment, microbial functional potentials, plant productivity, and ecosystem residual phosphorus. This initial model includes all biologically motivated pathways. Non-significant paths were subsequently removed, and the model was re-fitted to produce the reduced SEM shown in the main text (Figure 3).

**Figure S3.**
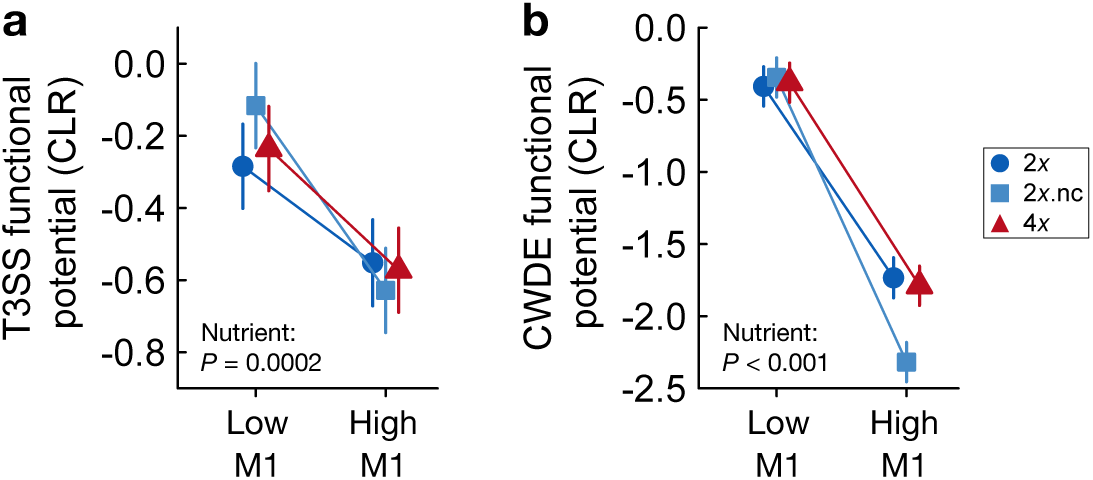
Pathogenesis-related functional potentials under nutrient enrichment. (a) Type III secretion system (T3SS) functional potential and (b) cell wall degrading enzyme (CWDE) functional potential across ploidy levels (2*x*, wild diploids; 2*x*.nc, treated but unconverted diploids; 4*x*, synthetic autotetraploids) and nutrient treatments (Low vs. High) in microbe-present microcosms (M1). Points show least-squares means (± SE) from general linear models (predictors: ploidy level, nutrient level, and their interaction; *n* = 143). Despite expectations that nutrient enrichment would increase pathogenicity, both T3SS and CWDE potentials were consistently higher under low nutrient supply across ploidy levels.

